# A metabolic perspective on eIF1A and eIF3B inhibition as a potential therapeutic strategy in psoriasis

**DOI:** 10.64898/2026.10.08.757342

**Authors:** Natalie Bordag, Benjamin Michel Rene Bourgeois, Tobias Madl, Peter Wolf, Nicole Golob-Schwarzl

**Affiliations:** Medical University of Graz, Department of Dermatology and Venereology, Graz, Austria; Otto Loewi Research Center, Division Medicinal Chemistry, Medical University of Graz, Graz, Austria; BioTechMed-Graz, Graz, Austria

## Abstract

Psoriasis is a chronic inflammatory skin disease marked by epidermal hyperplasia and distinct metabolic alterations. Here, we investigated the metabolic impact of targeting translation initiation factors by performing 1H NMR-based metabolomic profiling on polar skin extracts from imiquimod-induced (IMQ) and K5.TGFβ1 transgenic (TGFβ) mouse models treated with eIF1A or eIF3B siRNA.

IMQ-treated skin demonstrated significant, human-like metabolic shifts, including depleted energy substrates (glucose, glycogen, 3-hydroxybutyrate) and elevated phospholipid/nucleotide markers, which correlated strongly with epidermal thickness. In contrast, TGFβ skin exhibited no broad baseline polar metabolome changes. eIF1A/eIF3B inhibition selectively attenuated IMQ-associated energy and phospholipid alterations without causing widespread metabolic disruption in TGFβ skin.

Overall, eIF1A/eIF3B inhibition ameliorated psoriasis-like pathology and mitigated disease-associated metabolic demands in a context-dependent manner.

## Results and Discussion

Psoriasis is a chronic inflammatory skin disease characterised by epidermal hyperplasia, impaired keratinocyte differentiation, and immune activation. It is also associated with widespread local and systemic metabolic alterations. Reported changes include altered glucose utilisation and mitochondrial metabolism, phospholipid and fatty-acid remodelling, and markers of oxidative lipid damage, consistent with the metabolic demands of activated keratinocytes and immune cells [1]. Despite the availability of effective systemic biologics, additional topical treatment options are needed, especially for mild to moderate disease [2]. Translation initiation factors are emerging as therapeutic targets, particularly in cancer, although their central role in protein synthesis necessitates careful evaluation of their cellular effects [3]. We previously showed that topical eIF4E, eIF1A or eIF3B inhibition ameliorates psoriasis-like pathology in mouse and human 3D models [4,5].

Here, we investigated the metabolic effects of topical and systemic eIF1A or eIF3B siRNA inhibition by performing ^1^H NMR-based metabolomic profiling of polar dorsal skin extracts from imiquimod induced (IMQ) [6] or K5.TGFβ1 transgenic (TGFβ) [7] mice from our study [5]. Targeted and untargeted NMR features were log_10_-transformed and analysed using multivariate methods and generalised least-squares models (LOGGLS), and Pearson correlation analyses with epidermal thickness. Significance was set p_BH_<0.05 using Benjamini–Hochberg (BH) adjustment for multiple testing (see Supplementary Note and Data 1).

IMQ, but not TGFβ, was associated with a broad psoriasis-like polar metabolite profile (Fig. 1; Fig. S1). Consistent with these findings, broad alterations in metabolic and lipid pathway have been reported in IMQ-treated skin [8]. IMQ-treated skin showed significantly lower levels of glucose (contrast −0.64), glycogen (−0.78), 3-hydroxybutyrate (−0.91), and formate (−0.25) than controls. These changes are compatible with altered energy-substrate utilisation in highly proliferative and inflamed skin, align with the increased glucose consumption reported in human psoriasis lesions [1], and are consistent with metabolic adaptation to glucose limitation described in psoriatic keratinocytes [9]. Additionally, glycerophosphocholine (+0.65) and glycerol (+0.17) were significantly increased, indicating altered choline-associated phospholipid turnover or remodelling. Significantly increased arginine (+0.48) and uracil (+0.52), together with significantly reduced hypoxanthine (−0.25), suggested changes in amino acid and nucleotide metabolism, consistent with the biosynthetic demands of epidermal hyperplasia and immune activation.

**Figure 1:**
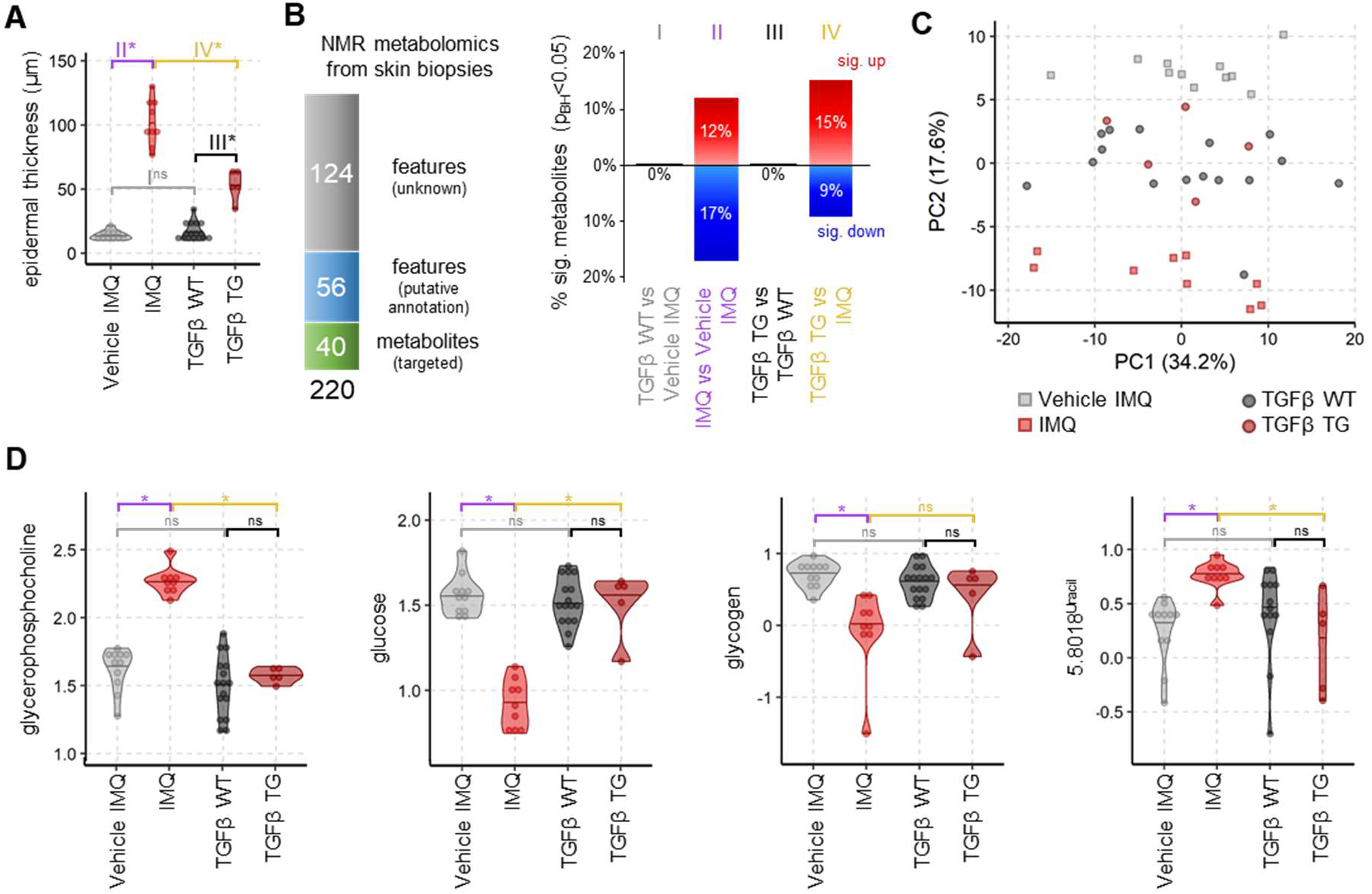
IMQ, but not TGFβ, is associated with marked skin metabolic remodeling. **A**, Epidermal thickness was significantly increased in IMQ and TGFβ mice compared to their respective controls. IMQ induced a more pronounced epidermal thickening than the TGFβ model, whereas control skin thickness did not differ between the two model systems. **B**, Overview of metabolite annotation and the proportion of significantly increased or decreased NMR features across the indicated pairwise comparisons (LOGGLS). IMQ induced widespread metabolic changes while no significant differences were detected for TGFβ mice relative to their respective controls. **C**, Principal component analysis (PCA) of 220 skin NMR features. Each point represents one mouse and proximity indicates similarity of metabolic profiles. The IMQ metabolomes differed distinctly from those of all other groups, with the groups separating along PC2. **D**, Selected metabolites consistent with altered glucose availability and phospholipid-associated metabolism in IMQ. Violin plots show log_10_(*peak area*); horizontal lines mark mean; *N = 5-16* per group. LOGGLS, Benjamini–Hochberg (BH) multiple test adjusted: ns = not significant, * p_BH_ < 0.05

In contrast, despite its established psoriasis-like epidermal and inflammatory phenotype, TGFβ-treated skin showed no significant differences in metabolite levels (Fig. 1; Fig. S1). Thus, the IMQ model recapitulated metabolic features implicated in human psoriatic disease, most notably altered glucose handling and choline/phospholipid-associated metabolism, whereas psoriasis-like pathology in TGFβ mice occurred without a broad shift in the polar skin metabolome.

We first assessed four individual regimens: topical eIF1A, systemic eIF1A, topical eIF3B, and systemic eIF3B inhibition. As differences between these regimens were limited and rare (<1% of NMR features reached statistical significance), all four treatment groups were pooled within each mouse model into a single treatment group (TRT) to reduce model complexity and increase statistical power.

Whereas 27% of detected features differed in IMQ skin, 36% of features were significantly different in TRT skin from controls (Fig. 2). The greater number of significant differences in TRT largely reflected the presence of additional features with small effect sizes rather than an exacerbation of IMQ-associated changes. In fact, many IMQ-associated changes were attenuated by TRT, although they often did not fully return to control levels (Fig. S2). For example, formate no longer differed significantly from the controls, while glucose, glycogen, 3-hydroxybutyrate, and glycerophosphocholine remained significantly altered, but showed smaller effect estimates than in untreated IMQ skin: glucose −0.64 in IMQ vs. −0.36 in TRT, glycogen −0.78 vs. −0.29, 3-hydroxybutyrate −0.91 vs. −0.69, and glycerophosphocholine +0.65 vs. +0.42. In contrast, significant changes in arginine, uracil, and hypoxanthine persisted at similar levels after TRT. Thus, eIF1A/eIF3B inhibition was associated with selective attenuation, rather than complete normalisation, of the IMQ-associated metabolic phenotype. This pattern is consistent with a reduction in inflammatory and proliferative metabolic demand accompanying improvement in psoriasis-like pathology.

**Figure 2:**
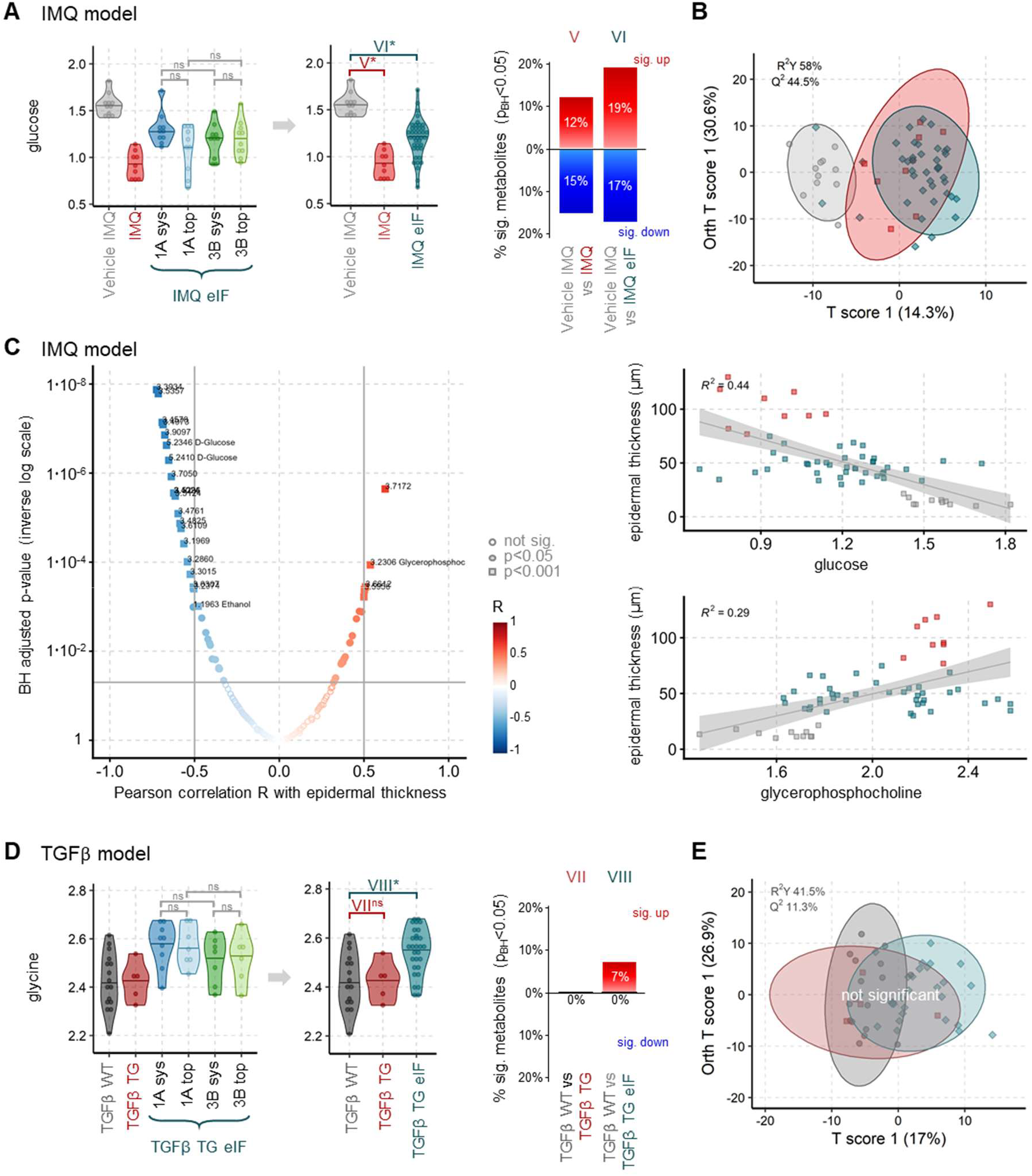
eIF1A or eIF3B inhibition attenuates IMQ-associated metabolic alterations, whereas the TGFβ skin metabolome remains largely stable. **A**, Representative glucose signal illustrating the rationale for pooling mice treated with topical or systemic eIF1A or eIF3B inhibition into a single treatment group (TRT). Differences among the four individual treatment regimens were small and rare (<1% of features significantly different by LOGGLS). The adjacent summary shows the proportion of significantly increased or decreased features in the indicated comparisons. **B**, Orthogonal projections to latent structures discriminant analysis (OPLS-DA) of controls, IMQ and TRT mice. Each point represents one mouse, proximity represents similarity, and ellipses indicate the group’s 95% confidence intervals. The model significantly distinguished the groups by 1,000-label permutation testing (p < 0.001). **C**, Each point represents one Pearson correlation between epidermal thickness and an individual NMR feature, with strong associations labelled (|R| > 0.5 and p_BH_ < 0.05). Among annotated metabolites, glucose was inversely associated with epidermal thickness, whereas glycerophosphocholine was positively associated with epidermal thickness. Regression lines and shaded areas indicate linear fits and 95% confidence intervals, respectively. **D**, Analogous to A, a representative glycine signal illustrating the rationale for pooling all treatments into one TRT group. The adjacent summary shows the proportion of significantly increased or decreased features in the indicated comparisons. **E**, OPLS-DA of controls, TGFβ and TRT mice showed no significant global separation by 1000-label permutation testing (p > 0.05), ellipses indicate the groups’ 95% confidence intervals. Violin plots show log_10_(*peak area*); horizontal lines mark mean; *N = 5-16* per group. LOGGLS, Benjamini– Hochberg (BH) multiple test adjusted: ns = not significant, ^*^ p_BH_ < 0.05

In contrast, TRT in TGFβ mice produced only a few small significant increases, with all contrast estimates ≤0.21, for example in glycine, alanine, serine, and valine (Fig. 2; Fig. S3). The limited metabolic response in treated TGFβ skin, compared with the partial attenuation of the extensive IMQ-associated profile, suggests a context-dependent metabolic effect of eIF1A/eIF3B inhibition rather than a general disruption of the detectable polar skin metabolome.

Pearson correlation analyses were performed separately within each mouse model, including control, diseased, and TRT samples, using epidermal thickness as a surrogate for disease severity (Fig. 2). Significant correlations were observed only in the IMQ model, while no significant correlations were found in TGFβ. In IMQ, epidermal thickness was negatively correlated with glucose (r = −0.62), 3-hydroxybutyrate (r = −0.51), glycogen (r = −0.43), and hypoxanthine (r = −0.39), whereas glycerophosphocholine (r = 0.42), uracil (r = 0.42), and glycerol (r = 0.35) were positively correlated. These associations mirror the disease-associated metabolic changes observed in IMQ skin and link alterations in energy-substrate handling, choline/phospholipid-associated metabolism, and nucleotide metabolism with epidermal thickening. As the analyses include controls, IMQ, and TRT, they demonstrate co-variation with disease severity but do not establish independent metabolic effects or causality.

Limitations include the incomplete coverage of the skin metabolome by ^1^H NMR analysis of polar extracts, particularly for complex lipids, as well as the analysis at a single endpoint, which precludes distinguishing direct metabolic effects of eIF inhibition from metabolic changes associated with reduced inflammation and epidermal hyperplasia. The absence of broad changes in TGFβ skin also does not exclude alterations beyond the detected polar metabolome or at other stages of disease. Nevertheless, the therapeutic improvement observed in both models indicates that eIF1A/eIF3B inhibition does not require reversal of a shared metabolic signature.

Psoriasis requires long-term treatment, making tolerability particularly important for therapeutic interventions. eIF1A/eIF3B inhibition improved psoriasis-like pathology in both models, attenuated IMQ-associated metabolic abnormalities, and did not cause broad detectable metabolic changes in TGFβ skin. These findings support further evaluation of translation-initiation targeting, including topical approaches, as a potential anti-psoriatic strategy that combines therapeutic activity with the potential to address inflammatory, proliferative, and metabolic disease features.

## Notes

Supplemental Information and Data associated with this article can be found in the online version.

Author contributions: Conceptualization, NGS, PW; Methodology, NGS; Software, NB, BMRB, TM; Validation, NGS, NB, PW; Formal analysis, NGS, NB, BMRB, TM; Investigation, NGS, NB, BMRB, TM; Resources, all authors; Data curation, all authors; Visualisation, NB; Supervision, NGS, TM, PW; Project administration, NGS, TM; Funding acquisition, NGS, BMRB, TM, PW; Writing – original draft, NB, Writing – Review & Editing, all authors;

We are very grateful for the technical assistance from Isabella Perchthaler, as well as to Sergio Oller-Moreno for the valuable discussions and advice. Austrian Science Fund FWF W1241 to PW, and the Cultural Office of the City of Graz and the Scientific Association of Styrian Dermatology supported the study. T.M. thanks the Center for Medical Research, Medical University of Graz, for laboratory access.

T.M. acknowledges the Austrian Science Fund (FWF) Excellence Cluster (grant DOI 10.55776/COE14) and FWF grants 10.55776/P28854 and 10.55776/I3792; Austrian Research Promotion Agency (FFG) grants 864690 and 870454; the Integrative Metabolism Research Center Graz; the Austrian Infrastructure Program 2016/2017; the Styrian Government (MetAGE, Zukunftsfonds, doc.fund program); the City of Graz (MetAGE, doc.fund); and BioTechMed-Graz (flagship project). This project was funded in part by the FFG and the European Union (EFRE) under grant 912192. For open access purposes, the author has applied a CC BY public copyright license to any author-accepted manuscript version arising from this submission.

The authors have declared that no conflict of interest exists. The authors used GPT-5.6 Terra strictly for language editing and accept full responsibility for all manuscript content

