## Supplementary Note for "A metabolic perspective on eIF1A and eIF3B inhibition as a potential therapeutic strategy in psoriasis"

### **Affiliations**

### **Supplementary Materials and Methods**

Skin samples analysed here were obtained from the same experimental cohorts described in our recent study of eIF1A and eIF3B inhibition in psoriasis-like skin inflammation [1].

#### **Animals**

All procedures were approved by the Austrian Federal Ministry of Education, Science and Research (BMWF-66.010/0032-II/3b/2013; 2021-0.393.968) and conducted according to institutional and national guidelines. Mice were housed at the Centre for Medical Research, Medical University of Graz, Austria, under either conventional or specific pathogen-free conditions, with ad libitum access to standard chow and water.

Psoriasis-like dermatitis was induced in female BALB/c mice (Charles River, Sulzfeld, Germany) by daily topical application of 62.5 mg of 5% imiquimod cream (MEDA Pharmaceuticals, Vienna, Austria) to the shaved dorsal skin for 7 consecutive days [2].

K5.TGFβ1 (TGFβ) transgenic mice, carrying full-length human TGFβ1 cDNA under the control of the keratin-5 promoter, were maintained through in-house breeding from breeding pairs provided by the model's creator, Prof. Xiao-Jing Wang [3].

#### ***In vivo* siRNA treatment**

As previously reported, the same siRNAs were used in cell culture, a human 3D psoriasis model, and mouse experiments, and target gene downregulation was confirmed by quantitative real-time PCR

(qRT-PCR) [1]. Transfection was performed using Metafectene®si+ transfection reagent (Biont, Munich, Germany) according to the manufacturer's instructions. The reported complete targeting sequences (5'–3' sense; DNA notation) were eIF1A/EIF1AX-C1, TTGGATAAATACCTCGGACAT; eIF1A/EIF1AX-C2, AAGCCTGTAAACTATGCATGA; eIF3B/EIF3B-C1, CGGGAAGATTGAACTCATCAA; and eIF3B/EIF3B-C2, GACCGACTTGAGAAACTCAAA. All siRNA were synthesized by QIAGEN (Hilden, Germany). Sequences had no overhangs, no chemical modifications and standard purification by manufacturer. A scrambled non-targeting control siRNA, UUCGUAGUACUCGAGUAGUC (QIAGEN, Germany) was used as control. The siRNA target sequences were highly conserved between human and mouse, with only minor nucleotide differences between the two species. For each target, C1 and C2 were administered an equimolar mixture.

Lyophilized 20 nmol of siRNA were reconstituted in 1 ml H<sub>2</sub>O. siRNA was administered by intraperitoneal injection or at a dose of 2 mg/kg body weight in a total volume of 200 µL per mouse. Immediately before use, siRNAs were diluted in phosphate-buffered saline (PBS) without a delivery reagent. Mice received target-specific or control siRNA once daily for 7 consecutive days by intraperitoneal injection (2 mg/kg body weight in 200 µL) or topical application to shaved dorsal skin (2 mg/kg in 200 µL). Topical treatment was gently spread over the skin surface 6 h after IMQ application. All siRNA treatments to all mice were administered at the same day time. Skin was collected 24 h after the final treatment.

#### **Nuclear magnetic resonance (NMR) sample preparation**

From each mouse 30–50 mg of skin tissue were resected and snap-frozen in liquid nitrogen, followed by storage at –80 °C until analysis. For metabolite extraction, 400 µL ice-cold methanol and 200 µL MilliQ H<sub>2</sub>O were added to each skin sample and transferred to 2 mL tubes with O-ring caps containing Precellys beads (1.4 mm zirconium oxide beads, Bertin Technologies, Villeurbanne, France) for homogenization by Precellys 24 tissue homogenizer (Bertin Technologies, Montigny-le Bretonneux, France). After centrifugation at 13,000 rpm for 45 min (4 °C), the supernatant was transferred to new 1.5 mL tubes and subsequently lyophilized at <1 Torr, 850 rpm, 25 °C for 10 h in a vacuum-drying chamber (Savant Speedvac SPD210 vacuum concentrator) with an attached cooling trap (Savant RVT450 refrigerated vapor trap) and vacuum pump (VLP120) (all Thermo Scientific, Waltham, MA, USA). The samples were re-dissolved in 500 µL of NMR buffer (0.08 M Na<sub>2</sub>HPO<sub>4</sub>, 5 mM TSP (3-(trimethylsilyl) propionic acid-2,2,3,3-d<sub>4</sub> sodium salt), 0.04 (w/v)% NaN<sub>3</sub> in D<sub>2</sub>O, pH adjusted to 7.4 with 8 M HCl and 5 M NaOH). The supernatants were transferred to 5 mm NMR tubes for data acquisition.

#### **<sup>1</sup>H NMR metabolomics data acquisition and targeted analysis**

Metabolic-profiling analysis was conducted at 310 K using a 600 MHz Bruker Avance Neo NMR spectrometer (Bruker Biospin, Rheinstetten, Germany) equipped with a TXI 600S3 probe head. The Carr–Purcell–Meiboom–Gill (CPMG) pulse sequence was used to acquire $^1\text{H}$ 1D NMR spectra with a pre-saturation for water suppression (cpmgpr1d, 512 scans, 73,728 points in F1, 12019.230 Hz spectral width, 1024 transients, recycle delay 4 s). NMR spectra were processed as previously described [2] and imported into Matlab2014b. TSP was used as the internal standard for chemical-shift referencing (set to 0 ppm). Regions around the water, TSP, and methanol signals were excluded. From the 112 total spectra two were excluded for technical issues not delivering valid measurements (TGFβ 3B topical, TGFβ 1A systemic). The NMR spectra were aligned, and a probabilistic quotient normalization (PQN) was performed. Reduced spectra and normalized spectra were generated by Matlab2014b, among which the normalized spectra were used to quantify metabolites by signal integration, as previously described [3].

#### **$^1\text{H}$ NMR untargeted peak extraction**

For untargeted data extraction the AlpsNMR package from R was used [4] with all 110 valid spectra at the same time. The AlpsNMR workflow comprised spectral interpolation ($-0.7$ to $10$ ppm), exclusion of solvent regions (water, $4.5$–$5.1$ ppm; methanol, $3.33$–$3.39$ ppm), and baseline correction by asymmetric least squares ($\lambda = 6$; $p = 0.01$). Peaks were then detected ($n\text{DivRange} = 0.1$ ppm; $\text{SNR.Th} = 3$), and a reference spectrum was selected for peak alignment (Supplementary Data 1). Peak alignment was performed with a maximum shift tolerance of $0.0015$ ppm. Non-informative regions which stabilized the alignment, were excluded ($-0.7$ – $0.9$ ppm and $9.0$ – $10.0$ ppm) afterwards to avoid biasing spectral normalization. Spectra were normalized by probabilistic quotient normalisation (PQN). Finally peaks were again detected on these baseline corrected, aligned and normalized spectra and peak areas were integrated allowing a maximum $0.004$ ppm distance between peaks for one metabolite. Peak areas were scaled by $10^8$ for easier readable numbers and all features with more than 30% negative values which were introduced during baseline correction were excluded as unreliable features. Extracted peaks were subjected to automatic identification against HMDB [5] and the top three hits were summarized as putative annotations. Targeted metabolites were also extracted with AlpsNMR with the same boundaries as used with Matlab2014b and results from both R and MatLab2014b correlated very well with each other. All data can be found in the supplementary data table 1.

#### **Statistical analysis**

Data visualisation and statistical analysis were performed with R v4.2.1 [6] (using the packages AlpsNMR, car, colorspace, correlation, dendsort, dplyr, emmeans, ggforce, ggplot2, ggpmisc, ggrepel, lawstat, MetaboAnalstR, mixOmics, nlme, openxlsx, RColorBrewer, readxl, rstatix, stringr, tidyr, tibble) [4,7,8] and TIBCO Spotfire v12.1.0, TIBCO, Palo Alto, CA.

All reported p-values were adjusted for multiple testing according to Benjamini–Hochberg (BH) denoted as $p_{BH}$ (R function *stats::p.adjust*).

Distribution and shomocedasticity were investigated with Kolmogorov-Smirnov test and Brown-Forsythe Levene-type test, respectively ($p_{BH}$ Supplementary Data 1). Over 98% NMR features were sufficiently normal and homoscedastic without transformation and with $\log_{10}$-transformation. The $\log_{10}$-transformation notably improved scedasticity (increased p-value) and performance of multivariate methods and was used for further statistical analysis.

Principal component analysis (PCA) was performed centred and scaled to unit variance (z-scaled) with *mixOmics::pca()* on all $\log_{10}(\text{peak areas})$ metabolites with multilevel factor *Exp\_ID*. The dataset contained no zeros and < 2% missing values which were imputed automatically with the inbuilt non-linear iterative partial least squares algorithm (NIPALS) approach.

For orthogonal projections to latent structures discriminant analysis (OPLS-DA) missing values were imputed with *MetaboAnalystR::ImputeMissingVar(., method = "knn\_var")*, data was scaled and centred to unit variance (z-scaled) with *MetaboAnalystR::Normalization(..., "AutoNorm")* and models were calculated with *MetaboAnalystR::OPLSR.Anal(., reg = TRUE)*, using experimental group as the response variable and standard 7-fold cross-validation. Model stability was additionally verified with 1000 random label permutations by *MetaboAnalystR::OPLSDA.Permut(., num = 1000)*.

Pearson correlations were calculated for each metabolite ($\log_{10}(\text{peak areas})$) against epidermal thickness ($\mu\text{m}$) with *correlation::correlation()*.

For univariate analysis of significant changes within each metabolite linear models were fitted with *nlme::gls()* and with *nlme::lme()* with the random factor $\sim 1|\text{Exp\_ID}$ to investigate potential experimental run differences. Models with and without random factor were compared within each metabolite for lower AIC (Akaike information criterion; relative estimate of information loss), higher log-likelihood (goodness of fit), significance in log likelihood ratio test comparing quality of Q-Q plots, randomness in residuals and impact on t-ratios. In all models less than 10% of the metabolites profited from addition of the random factor and t-ratios of these metabolites were very similar to those from models without the random factor. In accordance with model parsimony only results from models without random effect are reported. As analyses were performed on $\log_{10}$-transformed peak areas, estimated group differences are reported on the log scale. For brevity, this approach is referred to as LOGGLS ($\log_{10}$-transformed linear models using generalized least squares).

### Supplementary Figures and Tables

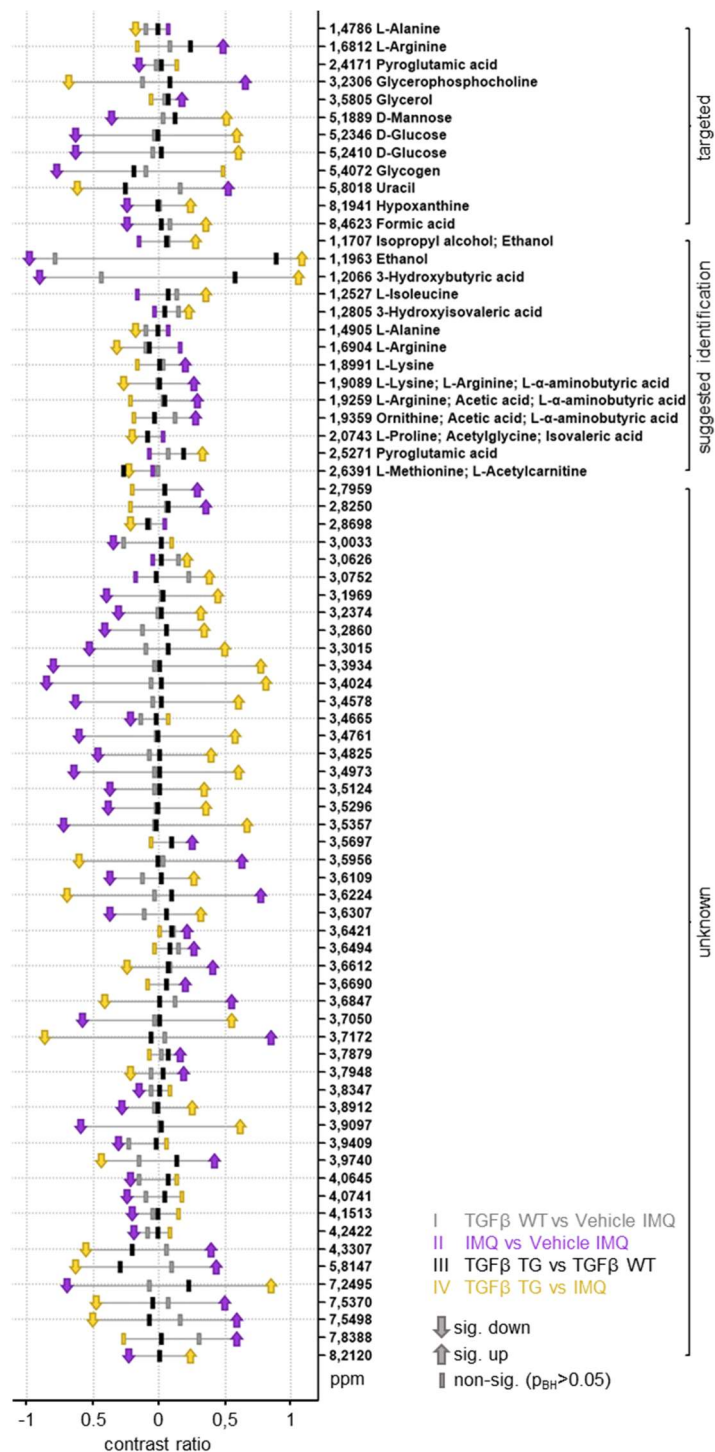

**Figure S1: IMQ induces widespread metabolic changes, whereas TGFβ mice show no significant global metabolic differences compared with WT controls. Dumbbell plot of LOGGLS results for metabolites that were significantly altered in at least one of the four comparisons (I–IV). The x-axis indicates the magnitude of metabolic changes, while statistical significance is indicated by symbol shape ($p_{BH} < 0.05$). Colors represent the four comparisons. $N = 5 - 16$ per group.**

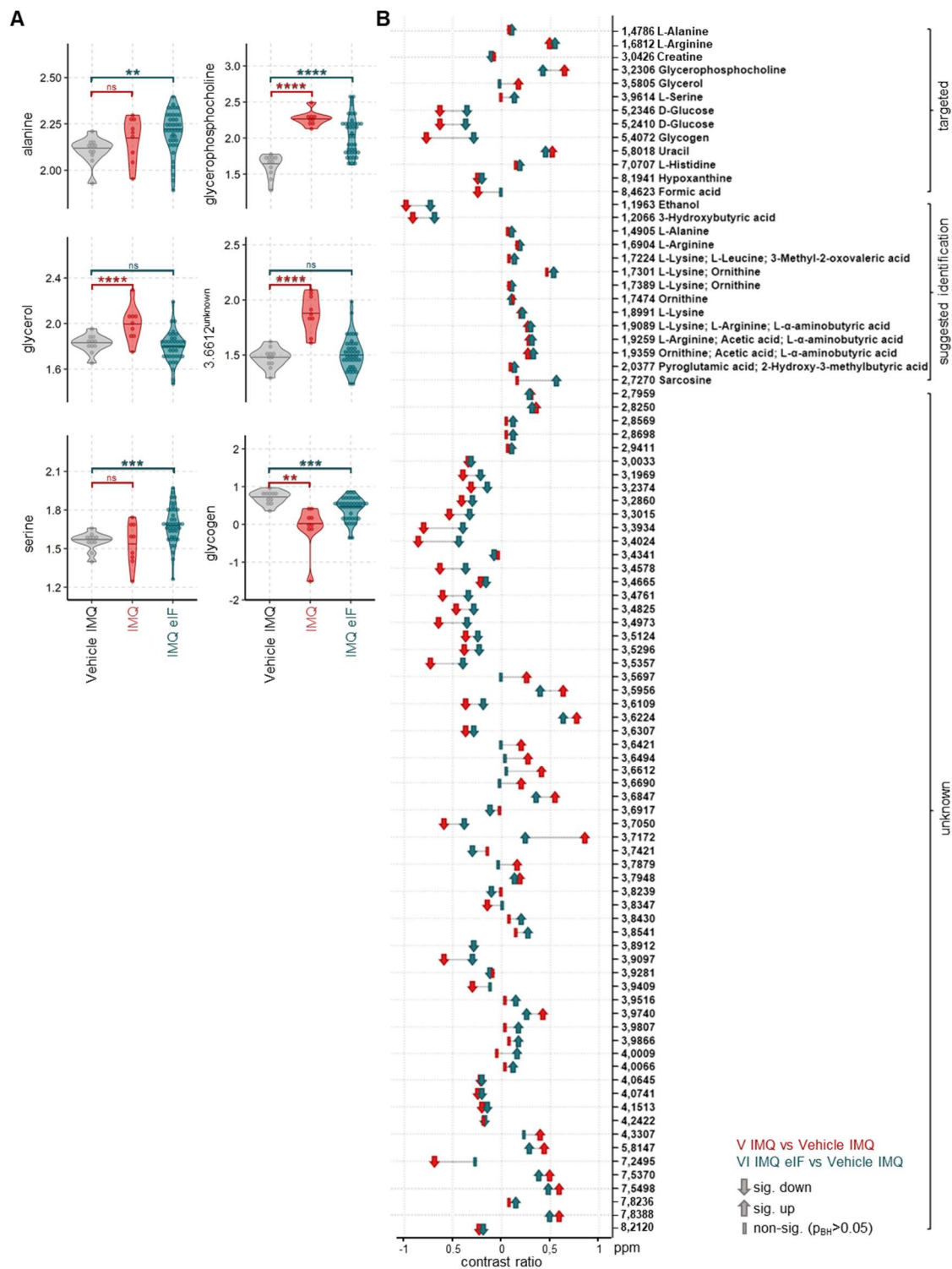

**Figure S2: eIF1A and eIF3B inhibition consistently ameliorates IMQ-induced metabolic changes.**

**A**, Violin plots showing the $\log_{10}(\text{peak area})$ of selected representative metabolites significantly altered in IMQ-treated mice, highlighting changes in glucose- and phospholipid-associated metabolic signals that are frequently ameliorated by treatment. **B**, Dumbbell plot of LOGGLS results for metabolites significantly altered in at least one of the two comparisons shown in A. The x-axis indicates the magnitude of metabolic changes, while statistical significance is indicated by symbol shape ($p_{BH} < 0.05$); colors represent the two comparisons. $N = 9 - 16$ per group. In all violin plots, horizontal lines indicate the mean. LOGGLS significances: ns = not significant. \* $p_{BH} < 0.05$; \*\* $p_{BH} < 0.01$; \*\*\* $p_{BH} < 0.001$; \*\*\*\* $p_{BH} < 0.0001$

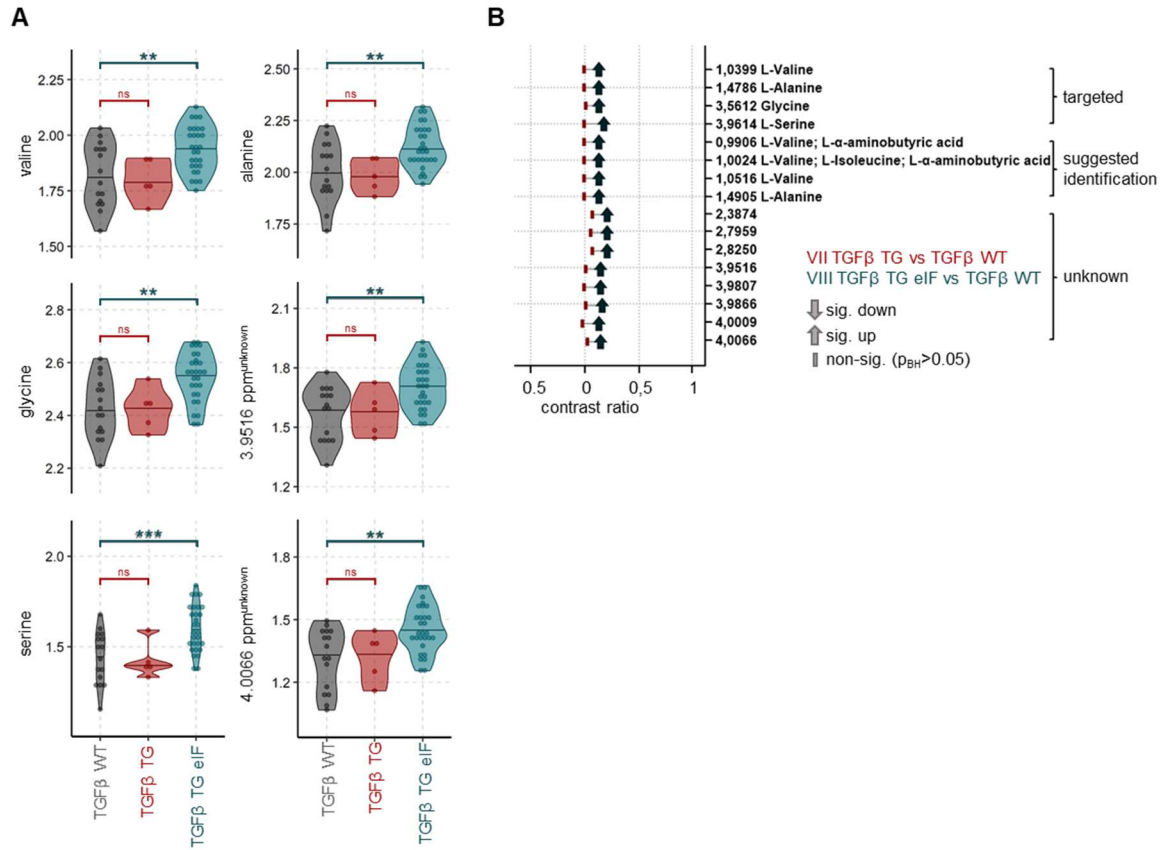

**Figure S3: Figure Sx: TGFβ expression and eIF1A or eIF3B inhibition have limited effects on the intradermal metabolome. A, Violin plots showing the log10(peak area) of selected representative metabolites that were significantly, but only weakly, increased in TGFβ-expressing mice following eIF1A or eIF3B inhibition. B, Dumbbell plot of LOGGLS results for metabolites significantly altered in at least one of the two comparisons. The x-axis indicates the magnitude of metabolic changes, while statistical significance is indicated by symbol shape ($p_{BH} < 0.05$); colors represent the two comparisons. $N = 9 - 16$ per group. In all violin plots, horizontal lines indicate the mean. $N = 9-16$ per group. In all Violin plots the horizontal lines mark the mean. LOGGLS significances: ns = not significant. \* $p_{BH} < 0.05$; \*\* $p_{BH} < 0.01$; \*\*\* $p_{BH} < 0.001$; \*\*\*\* $p_{BH} < 0.0001$**
